# Small-Angle and Quasi-Elastic Neutron Scattering from Polydisperse Oligolamellar Vesicles Containing Glycolipids

**DOI:** 10.1101/2024.11.26.625379

**Authors:** Lukas Bange, Amin Rahimzadeh, Tetiana Mukhina, Regine von Klitzing, Ingo Hoffmann, Emanuel Schneck

## Abstract

Glycolipids are known to stabilize biomembrane multilayers through preferential sugar-sugar interactions that act as weak transient membrane cross-links. Here, we use small-angle and quasi-elastic neutron scattering on oligolamellar phospholipid vesicles containing defined glycolipid fractions in order to elucidate the influence of glycolipids on membrane mechanics and dynamics. Small-angle neutron scattering (SANS) reveals that the oligolamellar vesicles (OLVs) obtained by extrusion are polydisperse with regard to the number of lamellae, *n*, which renders the interpretation of the quasi-elastic neutron spin echo (NSE) data non-trivial. To overcome this problem, we propose a method to model the NSE data in a rigorous fashion based on the obtained histograms of *n* and on their *q*-dependent intensity-weighted contribution. This procedure yields meaningful values for the bending rigidity of individual lipid membranes and insights into the mechanical coupling between adjacent membrane lamellae, including the effect of the glycolipids.

## Introduction

Naturally occurring membrane stacks, such as myelin sheaths or thylakoids (photosynthetic membranes), contain high amounts of glycolipids^1^ which are hypothesized to stabilize multilayered architectures.^2^ In fact, experiments with glycolipids^3^ and thylakoid lipid extracts^4^ have revealed spontaneous vesicle aggregation and membrane stack formation, and the hydration repulsion between glycolipid membranes was found to be of shorter range than that between zwitterionic phospholipids. ^5^ On the basis of an interplay between long-range van der Waals attraction and hydration repulsion, the adhesion energy between glycolipid bilayers was estimated to be enhanced by almost an order of magnitude with respect to commonly studied phosphatidylcholine (PC) lipid membranes.^2^ Within this picture, the tight cohesion between glycolipid membranes is rationalized solely on the basis of a shorter-ranged hydration repulsion. In other words, it does not explicitly invoke attraction between the saccharide groups belonging to the opposing membrane surfaces.

Other studies have, however, demonstrated that already small fractions of glycolipids can induce pronounced cohesion between lipid membranes,^6–8^ even against added electrostatic repulsion.^7–9^ Such favorable saccharide bonds had previously been considered to require specific sugar motifs like the Lewis^X^ trisaccharide.^6,7^ The finding that saccharide bonds are rather generic^10^ and occur for abundant sugar chemistries, ^8^ however, suggests that this phenomenon plays an important role in biology in light of the ubiquity of glycolipids in biological membranes. It is important to note that saccharide bonds are related to preferential saccharide-saccharide interactions^10^ and thus cannot be understood merely in terms of a weaker hydration repulsion. Moreover, the influence of such preferential interactions on the membrane separation can have different trends, if significant at all, depending on the size and chemistry of the saccharide headgroup.^7–9^

An aspect that has so far been neglected in the biological context is the effect that such saccharide bonds may have on the collective dynamics and mechanics of membranes. While these aspects are essential for the functionality of membranes, ^11^ little is known about the influence of saccharide bonds on the dynamics of membranes in biologically relevant multilayer configurations. Earlier studies using off-specular neutron scattering from oriented, solid-supported glycolipid membrane multilayers have been limited to the time-averaged structure of membrane stacks.^7,12^

The neutron spin-echo (NSE) technique is a powerful tool to investigate the dynamics of membrane undulations and the underlying mechanical properties.^13^ It is based on small wavelength changes as a result of quasi-elastic scattering and has been used successfully to investigate the influence of temperature,^14^ co-solutes, ^14,15^ as well as the addition of hard nanoparticles^16,17^ or peptides^18^ on membrane fluctuations. Gupta et al.^19^ give a comprehensive review. With very few exceptions^20^ the use of NSE for membrane investigations, however, has so far been limited to idealized model systems such as monodisperse small unilamellar vesicles (SUVs).^19^ But SUVs are intrinsically inappropriate for the study of membrane interactions because they do not feature any membranes in close proximity. On the other hand, oligolamellar vesicles (OLVs), which are much better suited, cannot always be produced such that they are monodisperse in size and in the number of lamellae. In fact, preparation protocols, for example based on extrusion through sub-micrometer pores, typically yield OLVs which are polydisperse in size and in the number of lamellae. In order to use NSE for such samples, new data analysis methods are thus required.

In the present work, we combine small-angle neutron scattering (SANS) and NSE to study OLVs formed by phospholipid membranes containing defined quantities of glycolipids with disaccharide headgroups. The SANS data provide detailed information on the distribution of lamellar numbers, which is essential for the meaningful interpretation of the NSE data. The analysis yields bending rigidities of individual membranes on the order of about 7 *k*_B_*T* and informs about the coupling strength between adjacent membranes. Our results suggest that the presence of glycolipids can indeed enhance membrane coupling.

## Results and discussion

The OLVs were based on the phospholipid 1-palmitoyl-2-oleoyl-glycero-3-phosphocholine (POPC) and additionally contained up to 20 mol% of a chain-unsaturated glycolipid, either Digalactosyl-diacylglycerol (DGDG) or N-hexadecanoyl-lactosyl-ceramide (LacCer), see Fig. 1 A. The low chain-melting phase transition temperature of POPC ensures that membranes are in the biologically relevant fluid phase at room temperature, where all measurements were conducted. As described in the methods section, the OLVs were prepared with extrusion pores of 400 nm diameter. For smaller pore sizes the resulting vesicles would be mostly unilamellar. For too large pores the resulting objects would be highly multilamellar and of ill-defined shape and size. After extrusion the OLVs exhibit polydispersity not only in the radius but also in the lamellar number *n*. The glycolipids are able to form saccharide bonds that hold adjacent membranes together, as schematically illustrated in Fig. 1 B. In earlier studies, this effect was found to be pronounced for LacCer lipids but much less pronounced for DGDG lipids.^8^

**Figure 1:**
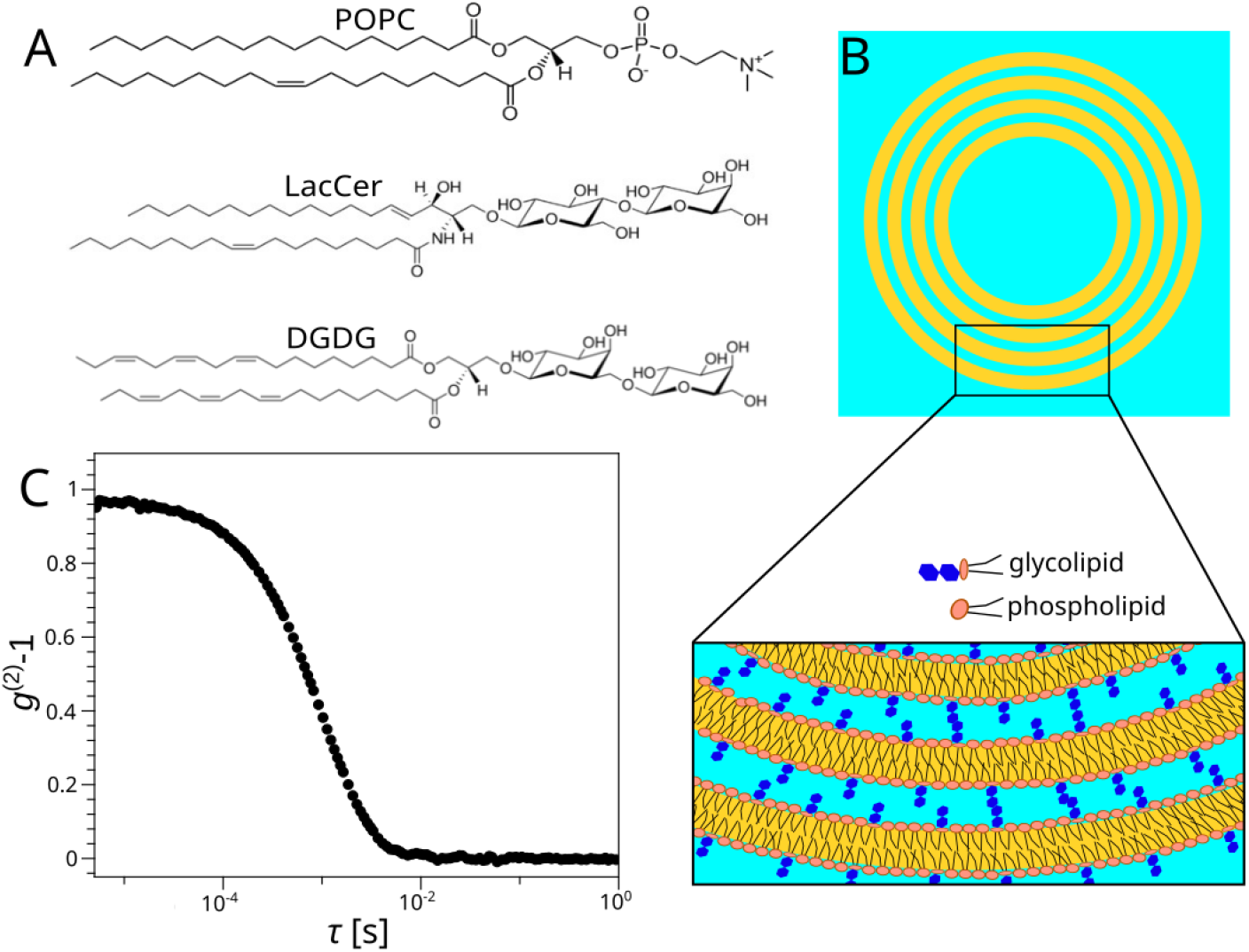
(A) Chemical structures of the lipids used: POPC, LacCer, and DGDG. The displayed structure of DGDG is the one with the most abundant hydrocarbon chain chemistry among a distribution. (B, top) Schematic representation of a sample composed of oligolamellar vesicles. (B, bottom) Zoom into the vesicles’ oligolamellar shell. Saccharide-saccharide bonds can form wherever disaccharides come into contact. (C) Representative DLS intensity correlation curve, *g*^(2)^(*τ*) − 1, obtained with OLVs composed of POPC with 20% LacCer.

### Basic characterization with DLS

At first, dynamic light scattering (DLS) measurements were carried out to determine the size distribution of the extruded OLVs. In DLS, a laser beam is scattered in the sample with the scattered light being measured as a function of time and scattering angle. From the temporal correlation of fluctuations in the scattered light intensity the size distribution of particles in terms of their hydrodynamic radii can be reconstructed.^21^ A typical correlation function, *g*^(2)^(*τ*) − 1, is shown in Fig. 1 B. The average hydrodynamic OLV radii 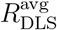, obtained as described in the methods section, are summarized in Table 1. The meaning of *R*_DLS_ is also illustrated in Fig. 3. With approximately 200-400 nm, the corresponding OLV diameters are comparable to the diameter of the extrusion pores. Note that this result is non-trivial because substantial differences between extrusion pore and vesicle diameters have been observed before. ^22^ We refrain from trying to rationalize the different sizes obtained with different lipid compositions, because the mechanisms of OLV formation through extrusion are complicated and influenced by many factors that may be affected by the lipid composition. DLS also confirmed the expectation of a significant polydispersity in the radius, with polydispersity indices (PDI) of about 0.1 - 0.15, which requires that 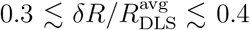, because 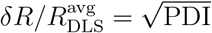.

**Table 1:**
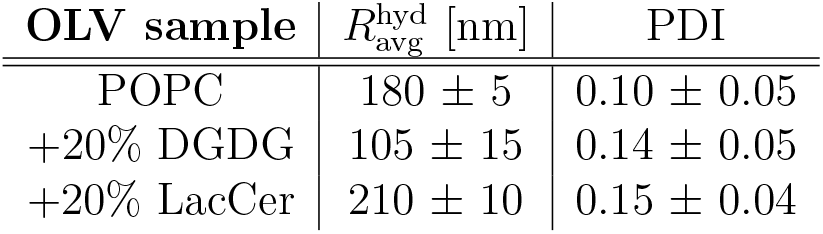
Sample characteristics as determined by DLS. Errors represent the standard deviation of measurements at all angles.

| OLV sample | $R_{\text{avg}}^{\text{hyd}}$ [nm] | PDI |
| --- | --- | --- |
| POPC | $180 \pm 5$ | $0.10 \pm 0.05$ |
| +20% DGDG | $105 \pm 15$ | $0.14 \pm 0.05$ |
| +20% LacCer | $210 \pm 10$ | $0.15 \pm 0.04$ |

### Structural characterization with SANS

In the next step, the OLVs were characterized by SANS in order to determine their structural characteristics. As will be discussed further below, this information is essential for the interpretation of the NSE data. Fig. 2 A shows a representative SANS curve, *I*(*q*), obtained with POPC OLVs containing 20% DGDG. The first qualitative conclusions can be drawn directly without any model-based data analysis. Most importantly, a distinct intensity peak is visible at *q* ≈ 0.1 Å^−1^. This value corresponds to a periodic spacing of *d* = 2*π/q* ≈ 6 nm, which is typical for lamellar phases of phospholipid bilayers^23^ including those consisting purely of POPC^24^ and coincides with the combined thickness ^25^ of the bilayer (*d*_B_) and the interstitial water layer (*d*_W_), *d* = *d*_B_ + *d*_W_. For small vesicles which are reported to be unilamellar, this peak is absent.^24,26^ Table 2 summarizes the *d* values obtained from the peak positions in all samples investigated. The spacing in the OLVs of pure POPC (*d* = 6.2 nm) is in satisfactory agreement with values of ≈ 6.3 nm reported earlier. ^27,28^ Samples containing glycolipids have very similar spacing and appear to exhibit a slight trend towards lower spacing (*d* = 6.0 - 6.1 nm), in line with earlier reports for glycolipids with the same disaccharide headgroups.^8,9^ These weak trends have to do with an interplay between the saccharides’ preferential binding modes and steric interactions^29^ and we are not going to further interpret them here.

**Table 2:** Characteristics of the different OLV samples as determined by SANS. See Methods section for details on the error estimates.

| OLV sample | $d$ [nm]<br>($\pm 0.05$) | $\mu$<br>($\pm 0.1$) | $\xi$<br>($\pm 0.2$) | $\nu$<br>($\pm 0.1$) | $n_{\text{avg}}$<br>($\pm 0.2$) | $R_{\text{SANS}}^{\text{avg}}$ [nm] | $\delta R/R_{\text{SANS}}^{\text{avg}}$ |
| --- | --- | --- | --- | --- | --- | --- | --- |
| POPC | 6.2 | 0.9 | 0.7 | 0.7 | 2.0 | $\gtrsim 100$ | $\gtrsim 0.15$ |
| +10% DGDG | 6.1 | 1.0 | 0.4 | 0.5 | 1.7 | $\gtrsim 100$ | $\gtrsim 0.15$ |
| +20% DGDG | 6.0 | 1.0 | 0.4 | 0.5 | 1.7 | $\gtrsim 100$ | $\gtrsim 0.15$ |
| +20% LacCer | 6.1 | 1.0 | 0.8 | 0.6 | 2.5 | $\gtrsim 100$ | $\gtrsim 0.15$ |

**Figure 2:**
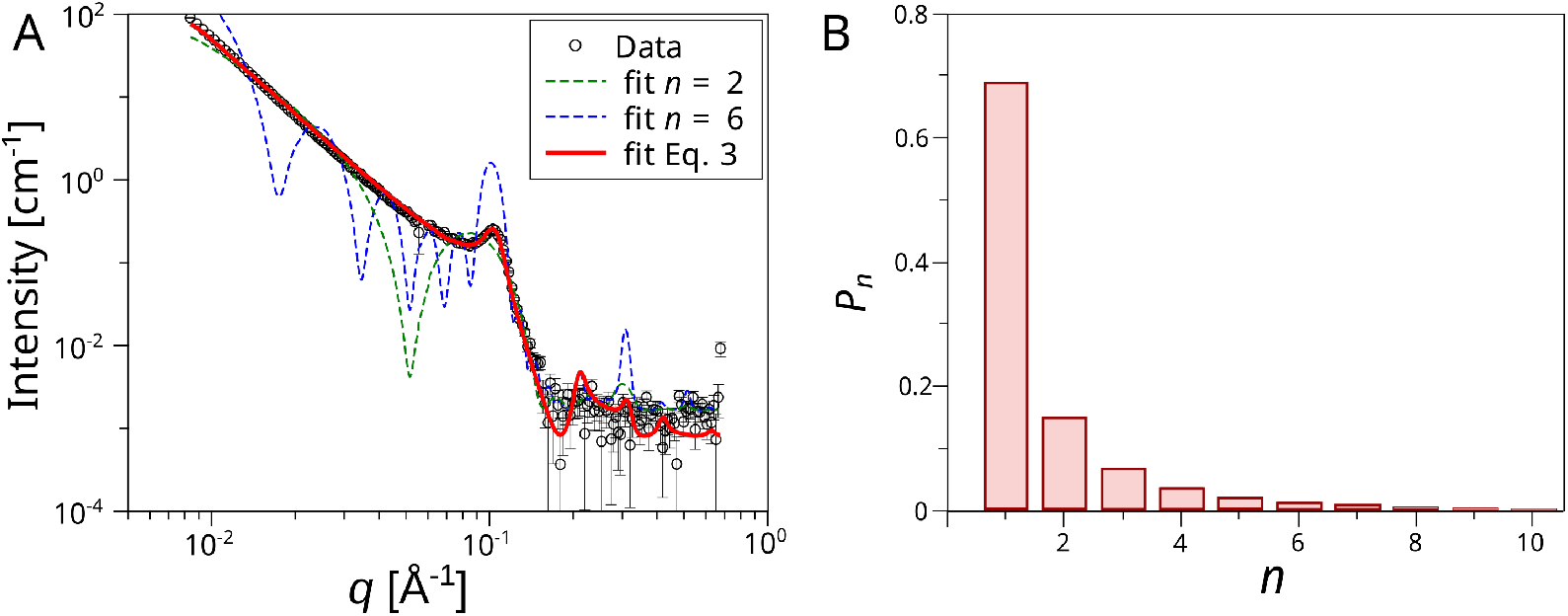
(A) Representative SANS curve after background subtraction, obtained with POPC OLVs containing 20% DGDG. Lines indicate attempts to fit the data. Fits indicated with dashed lines assume monodispersity in the number *n* of lamellae; the fit indicated with the solid line assumes a distribution of *n* (Eqs. 1 and 2), which is shown in panel (B).

**Figure 3:**
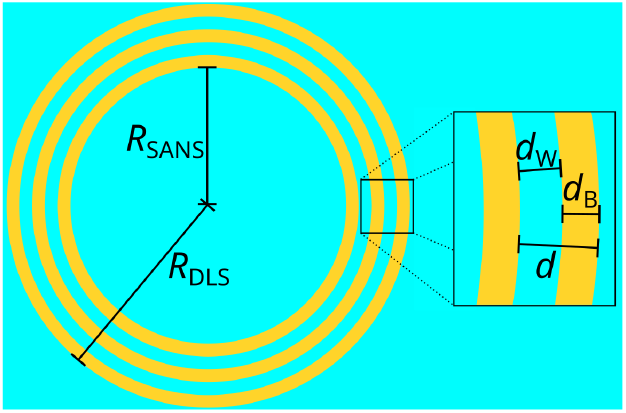
Schematic illustration of an OLV with lamellar period *d*, bilayer thickness *d*_B_, water layer thickness *d*_W_, and with inner and outer vesicle radii *R*_SANS_ and *R*_DLS_, respectively. In the shown example, the number of lamellae is *n* = 3.

As seen in Fig. 2 A, the peaks are not very pronounced, which confirms that the vesicles are oligolamellar. Regarding the size of the OLVs, the absence of a plateau at the lower end of the *q*-range requires that the OLV radius is larger than ≈ 100 nm, which is consistent with the DLS results presented before, irrespective of the slightly different definition of the radius in the two methods (see Fig. 3, recalling that *R*_DLS_ ≈ *R*_SANS_ *>> nd*, where *n* is the number of lamellae). Moreover, the absence of other low-*q* features requires that the OLVs exhibit substantial polydispersity with regard to their radius 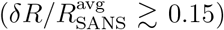, which is again consistent with the DLS results (see previous subsection).

Having established the general sample architecture, a model to describe the SANS data can be formulated as illustrated in Fig. 3. In the model, the OLVs are assumed to have spherical shapes with average inner radius 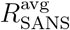, noting that the SANS signal in the covered *q*-range is vastly insensitive to the exact outer shape, as long as the OLVs are large enough. To account for polydispersity in the radius, the model averages over a large number of OLVs, each with its individual radius *R*_SANS_ (see Fig. 3), which is randomly picked from a normal distribution of width *δR* around 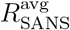. To account for the oligolamellarity, each OLV is described as a set of concentric membrane shells, the number of which (i.e., the lamellar number) is termed *n*. The center-to-center distance between two consecutive membrane shells coincides with the lamellar period *d* = *d*_B_ + *d*_W_ introduced before. Unfortunately, due to limited statistics at high *q*, the thickness parameter *d* cannot reliably decomposed into the thickness of the water layer *d*_W_ and the thickness of the bilayer *d*_B_. Therefore for *d*_B_ we resort to the well-established literature values for such lipids (*d*_B_ = 32 Å^23,25^), noting that the analysis is robust with respect to the precise value. The water layer thickness *d*_W_ then follows accordingly for a given choice of *d*. The neutron scattering length density (SLD) of the D_2_O surrounding the bilayers is *ρ*_W_ = 6.37 · 10^−6^ Å^−2^. The SLD of the bilayers can be calculated from the SLDs and volumes of the headgroups and hydrocarbon chains, ^30^ which yields an approximately vanishing SLD (*ρ*_B_ ≈ 0). The SANS curve for each individual parameterized OLV is calculated as described in the methods section. The SLD difference between bilayers and the surrounding D_2_O matters for the overall intensity scale. In light of the substantial polydispersity in the OLV characteristics and for the sake of simplicity, the absolute intensity of the modeled signals is, however, adjusted by treating the overall lipid volume fraction as an adjustable parameter. The obtained volume fraction is systematically lower (e.g., 0.31 vol% for the data in Fig. 2 A) than the nominal one (0.48 vol%), which can be attributed to uncertainties in the weighed amount, in the estimated SLD difference, and to sedimentation losses upon transfers between containers. As shown in the Supporting Information (Fig. S5), the SANS data cannot be reproduced when imposing the nominal lipid volume fraction, even when readjusting the other model parameters.

At first we made attempts to model the SANS data under the assumption that all OLVs have the same number *n* of lamellae, in which case one is left with six independent adjustable parameters: 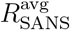, *δR, d*, and *n*, as well as the lipid volume fraction and a parameter accounting for a flat incoherent background. The dashed lines in Fig. 2 A are theoretical SANS curves corresponding to these attempts. While the dashed blue line (assuming *n* = 6) reproduces the width of the peak but fails to reproduce the peak height and the featureless monotonic intensity decay at lower *q*, the green line (assuming *n* = 2) roughly reproduces the peak height and the low-*q* region but fails with regard to the peak width. As it turns out, it is not possible to satisfactorily reproduce the entire SANS curve with a model that assumes the same *n* for all vesicles.

In the next step, we therefore allow for a probability distribution *P*_*n*_ of lamellar numbers *n*. To this end, polydispersity in *n* between different OLVs is formally considered, although, in practice, allowing for polydispersity within individual OLVs would lead to consistent results. The theoretical SANS curve *I*(*q*) is described as the weighted superposition of the contributions *I*_*n*_(*q*) originating from different *n*:

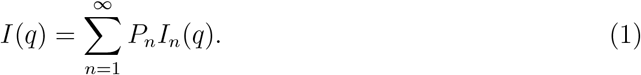

In order to model *P*_*n*_, we use a stretched exponential function for convenience, because this function covers a wide spectrum of shapes with only few adjustable parameters:

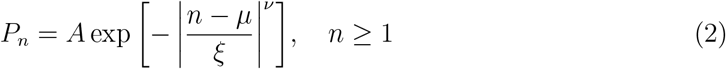

where *ξ* is the decay length, ν is the stretching exponent, *µ* defines where the distribution has its maximum, and the pre-factor *A* must fulfill the normalization condition 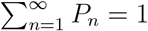. The average lamellar number for each set of parameters follows as

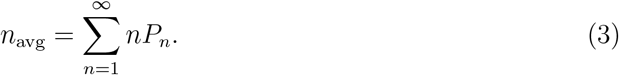

In total, the model then has eight independent adjustable parameters: 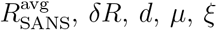, *δR, d, µ, ξ*, and ν, as well as the two parameters for scaling and background. The solid line in Fig. 2 A is the theoretical SANS curve corresponding to the best-matching set of parameter values. It is seen that this model, which allows for a distribution of *n*, reproduces the experimental data well, which confirms the suitability of the functional form assumed in Eq. 2. As shown in the Supporting Information (Fig. S1), the same is the case for the experimental data of the other samples. The distribution *P*_*n*_ corresponding to the theoretical model (red line) in Fig. 2 A is presented in Fig. 2 B and characterized by *µ* ≈ 1.0, *ξ* ≈ 0.4, and ν ≈ 0.5, which yields *n*_avg_ ≈ 1.7 according to Eq. 3. Table 2 summarizes the best-matching parameters for all samples investigated. The fits require that 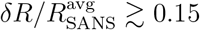, in line with the DLS results, and we generally find that *n*_avg_ ≈ 2 and that the distributions are rather broad. There are no obvious systematic trends with regard to the influence of the glycolipids on the sample characteristics, however the highest average number of lamellae is found for POPC with 20 mol% LacCer, which points towards some enhanced cohesion between the membranes. In summary, we find that the OLVs prepared by the extrusion procedure exhibit considerable polydispersity in both *R*_SANS_ and *n*. This structural complexity poses a challenge to the NSE data analysis, because each vesicle (or, equivalently, each vesicle portion with its local *n*) thus has a different *q*-dependent contribution to the measured NSE signal. However, as will be shown in the next section, these contributions can be quantified with the SANS-based parameters in Table 2.

### Membrane dynamics and mechanics as obtained by NSE

Fig. 4 shows a representative set of NSE data, where the intermediate scattering function *S*(*q, t*) is plotted for various *q*-values. The intensity systematically decays with *t*, the faster the higher *q* is. The decay is not a simple exponential, which is easily seen when plotting log(*S*) vs. *t* (see Supporting Information Fig. S2). Qualitatively, such behavior is expected for thermally fluctuating layers like lipid membranes and is typically described within the framework of the Zilman-Granek model: ^13^

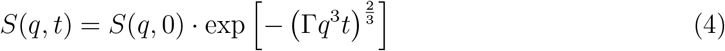

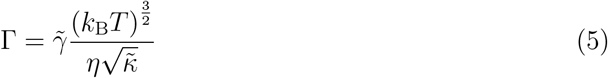

where *η* is the solvent viscosity, *k*_B_*T* the thermal energy, 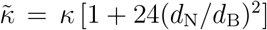 is the renormalized layer bending rigidity,^31,32^ *κ* is the regular bending rigidity, *d*_N_ is the distance of the bending-neutral surface in a half-layer from the center of the layer and *d*_B_ is the bilayer thickness.^33^ Assuming a fixed ratio *d*_N_*/d*_B_, this simply translates to a factor between *κ* and 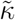. The nominal value of the dimension-less pre-factor 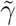 is 0.025.^13^ For the typical assumption regarding *d*_N_ and by accordingly replacing 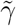 with *γ* = 0.0069, one obtains a simpler yet equivalent expression for Γ in terms of the regular bending rigidity alone: ^34^

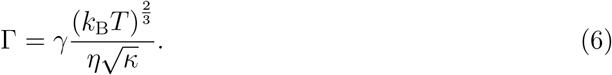

**Figure 4:**
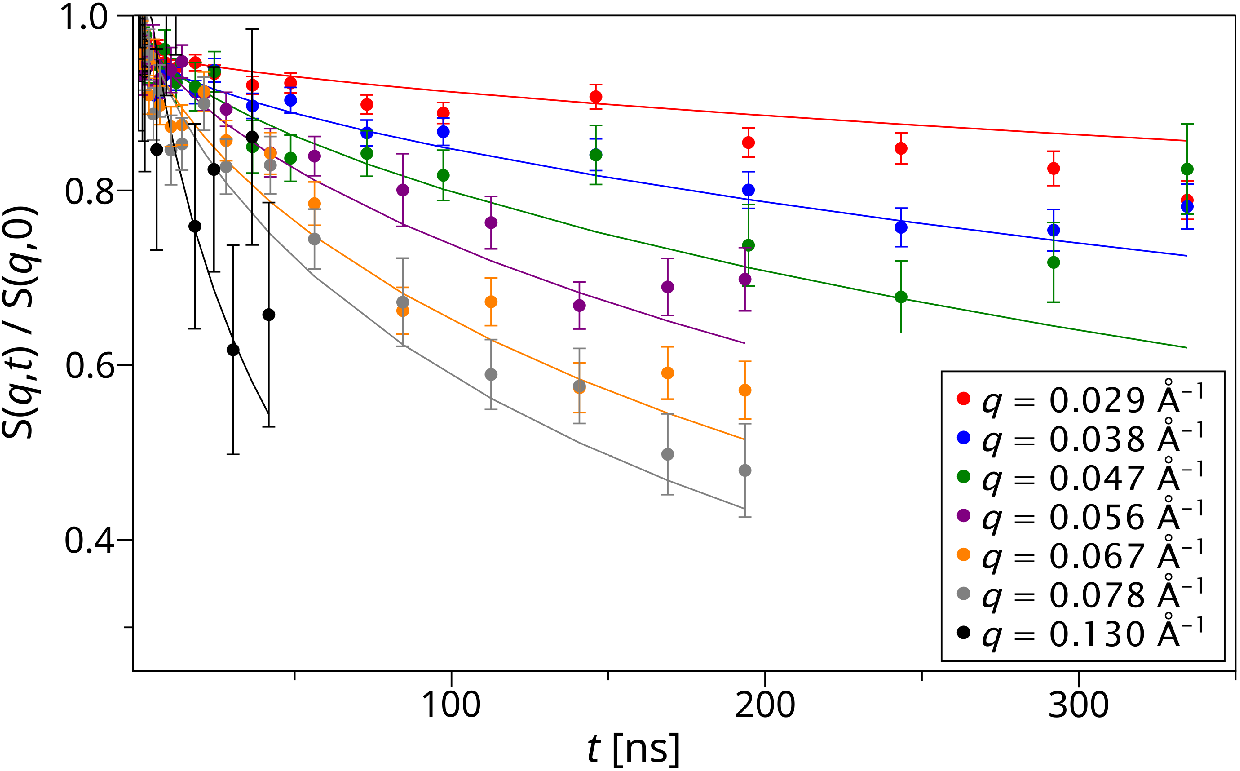
Normalized intermediate scattering function determined by NSE for POPC OLVs containing 20% DGDG. Solid lines indicate the modeled intensities based on Eq. 10 for the best matching parameters *κ*_1_ and *α* (see Table 3).

Eqs. 4 to 6 take into account neither the translational diffusion of the vesicles nor effects from their finite size and spherical shape. In a recent paper, Granek et al. ^35^ published an expression which explicitly does so. However, for relatively large vesicles with diameters over 200 nm, such as the ones investigated here, the effects are rather small and we are mainly interested in relative changes of *κ*, which is why we use the simple stretched exponential form of the Zilman-Granek theory. In the idealized case of a single membrane, the curves for all *q*-values are characterized by the same value of Γ. Since 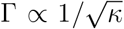, lower Γ values are indicative of higher effective layer rigidities. As shown in the Supporting Information (Fig. S3), the Γ values of OLVs loaded with glycolipids are systematically somewhat lower than those of the pure POPC OLVs. While this observation appears to be a qualitative indication for some glycolipid-induced rigidification, we remark that the effective rigidity of oligolamellar membranes is influenced by several factors, notably the rigidity of a single membrane, denoted as *κ*_1_ in the following, the number *n* of membranes, and the extent of inter-membrane coupling. As seen in Table 2, there are considerable variations in *n*_avg_, which affect Γ and therefore obscure the influence of the glycolipids on membrane coupling and bending rigidities. A more involved approach to model the data thus has to be taken.

**Table 3:** Single-membrane bending rigidity *κ*_1_ and coupling exponent *α* as determined by NSE. Error estimates correspond to the one-sigma confidence interval.

| OLV sample | $\kappa_1 [k_B T] (\pm 1.0)$ | $\alpha$ |
| --- | --- | --- |
| POPC | 6.5 | 1.0 (fixed) |
|  | 5.1 | 2.0 (fixed) |
| +10% DGDG | 11.7 | 1.0 (fixed) |
|  | 9.5 | 2.0 (fixed) |
| | 6.5 (fixed) | $2.0 \pm 0.3$ |
| +20% DGDG | 12.2 | 1.0 (fixed) |
|  | 10.1 | 2.0 (fixed) |
| | 6.5 (fixed) | $2.2 \pm 0.3$ |
| +20% LacCer | 31.3 | 1.0 (fixed) |
|  | 20.8 | 2.0 (fixed) |
|  | 15.2 | 3.0 (fixed) |
| | 6.5 (fixed) | $4.8 \pm 0.5$ |

At first we note that, when OLVs with different *n* coexist, their relative contributions to the intermediate scattering function depend on *q*. This is because *S*(*q, t*) is intensity-weighted and the intensity for each *n* has a different *q*-dependence (see *I*_*n*_(*q*) in Eq. 1). In fact, as shown in the Supporting Information (Fig. S4), one obtains different Γ values when using Eq. 4 for fits at different *q*. However, with the SANS intensities *I*_*n*_(*q*) at hand we can readily work out the *q*-dependent relative intensity contributions *c*_*n*_(*q*) of each lamellar number *n*:

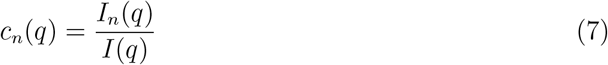

which by construction fulfil the requirement

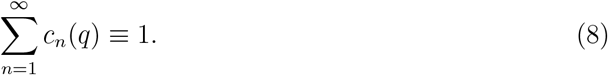

Fig. 5 shows a set of *c*_*n*_(*q*) for a typical parameter set. The *q*-values for which NSE measurements were carried out are indicated with vertical lines. It is seen that different *n* become dominant at different *q*.

**Figure 5:**
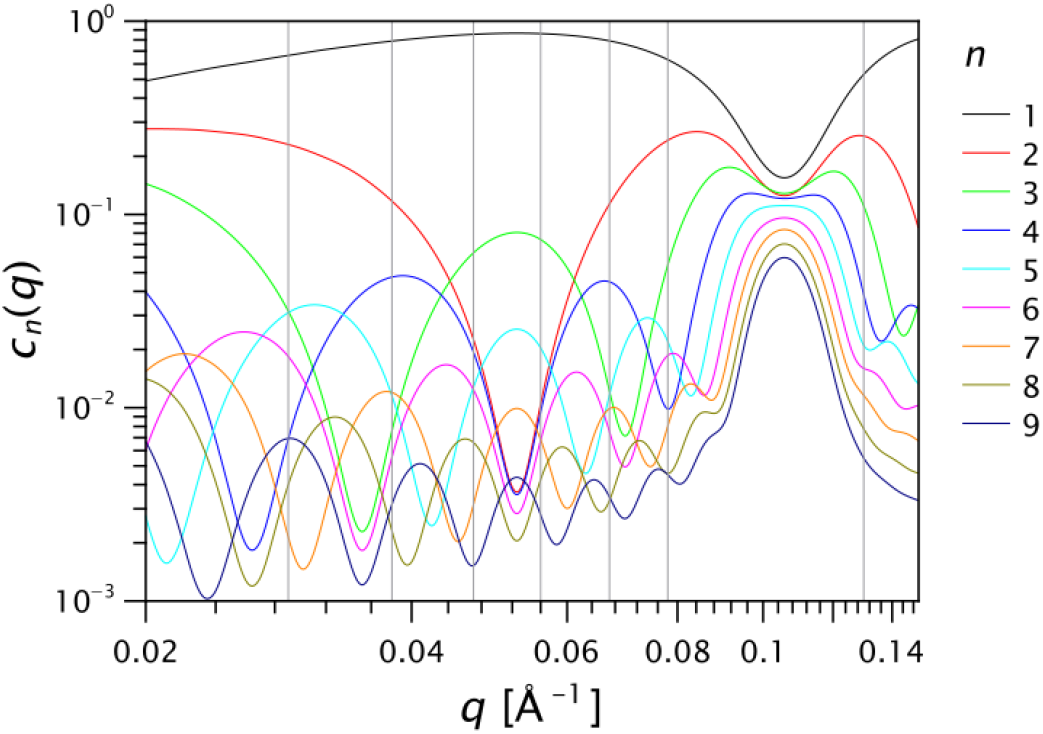
Relative intensity contributions *c*_*n*_ of different lamellar numbers *n* as a function of *q* for the parameter set specified in Table 2 for POPC OLVs containing 20% DGDG. Vertical lines indicate the *q*-values specified in Fig. 4 at which NSE was measured.

In the next step we assume that the bending rigidity *κ*_*n*_ of a stack of *n* membranes has the following functional dependence on the bending rigidity *κ*_1_ of a single membrane:

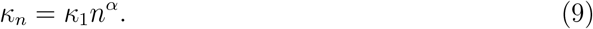

The coupling exponent *α* ∈ [1, 3] can assume any value between the limit of fully uncoupled membranes that are deformed independently (*α* = 1) and the extreme case of fully coupled membranes that are deformed as a single rigid object whose thickness is the entire oligolamellar layer (*α* = 3, because the bending stiffness then scales with the third power of thickness according to the well-known beam equation). This description can be understood as a generalization with respect to an earlier work^36^ in which a quadratic dependence of *κ* on *n* was found, i.e., *α* = 2, in line with the predictions by Evans^37^ and by Deserno^38^ for the case of a single fluid membrane.

Finally, by combining Eqs. 7-9 with Eqs. 4 and 6 we obtain an expression for the intermediate scattering function of polydisperse OLVs with mechanically coupled membranes:

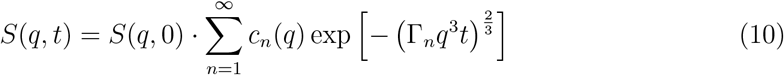

where

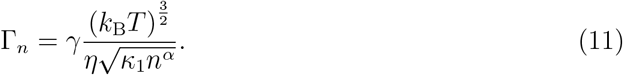

Since the coefficients *c*_*n*_(*q*) are already known from the preceding SANS analysis (see Fig. 5), we are left with only two unknown parameters, *κ*_1_ and *α*, which can both be adjusted to match the experimental NSE data. In the fitting procedure, *q*-values in the Bragg peak are excluded^39^ in order to avoid artifacts due to de Gennes narrowing, which results in a slowing down of the dynamics near the peak position.^40^

As exemplified in Fig. 4, Eq. 10 simultaneously reproduces the complete set of NSE data measured at various *q* (see Supporting Information Figs. S6 for all data sets). The obtained parameters are summarized in Table 3. Irrespective of the choice of *α*, the obtained single-membrane bending rigidity *κ*_1_ of POPC is of the order of 10 *k*_B_*T*, which is in convincing agreement with earlier studies using NSE on SUVs, ^19,41,42^ off-specular neutron scattering, ^7,12^ or other experimental techniques.^43^ For pure POPC membranes one may expect *α* ≈ 1 because PC bilayers are known to slide along each other with little friction. ^44^ At this point it should however be noted that, when a limited *q*-range is sampled with only a handful of data points, there is a considerable covariance between *κ*_1_ and *α* in the fit, such that these two parameters cannot be determined independently in a reliable manner. We therefore test two limiting cases as hypotheses.

At first, we assume that the presence of glycolipids does not affect coupling between membranes. In this case reproducing the experimental data requires a substantial increase in *κ*_1_ when glycolipids are added to the membranes. For example, when *α* is fixed to 1, *κ*_1_ increases from ≈ 7 *k*_B_*T* for pure POPC to ≳ 30 *k*_B_*T* for POPC with 20 mol% LacCer. A similar trend is obtained when considerable coupling is assumed and *α* is fixed to 2, see Table 3. But it is rather implausible that the bending rigidity of a lipid bilayer in the fluid phase increases so much by the addition of 20% glycolipids. In fact, as measured and rationalized earlier, ^7^ the effect of even rather bulky saccharide headgroups on the bending rigidity is negligible as long as they are not densely packed laterally. A clear influence of the saccharide headgroups was reported only at high densities (pure glycolipid without matrix). ^12,45^ Furthermore, the glycolipids used here have a higher degree of chain unsaturation than the POPC matrix lipid, which has a rigidity-reducing effect according to earlier reports. ^33^ It is therefore unlikely that *κ*_1_ increases when the glycolipids are introduced. We therefore assume in the next step that the presence of glycolipids affects coupling between membranes but leaves the bending rigidity unaffected and equal to the value obtained with pure POPC. In this case reproducing the experimental data requires a substantial increase in *α* when glycolipids are added to the membranes. For example, for *κ*_1_ = 6.5 *k*_B_*T* (see Table 3), the coupling exponent increases from *α* = 1 for pure POPC to *α* ≈ 2 for POPC with 10 or 20 mol% DGDG. For POPC with 20 mo% LacCer *α* reaches an even higher value. Such an increase in the coupling is qualitatively expected when saccharide bonds are formed between glycolipids of the opposing membranes. A coupling constant exceeding 2 indicates that the saccharide bonds do not only constrain the distance between adjacent membranes, but also constrain the lipids’ lateral movement. Namely, the bonds can resist to the relative sliding movement of adjacent membranes at least transiently because the two binding partners are laterally co-localized and have fixed positions in the membranes on time scales shorter than those of the relevant lipid lateral mobility. Noteworthy, the increase in *α* is most pronounced for the sample containing the LacCer headgroup, which was previously found to cross-link membranes more strongly than glycolipids with DGDG headgroups. ^8^ In this case, when fixing *κ*_1_ at 6.5 *k*_B_*T*, the value obtained for *α* even exceeds the upper plausible limit of 3, indicating that *κ*_1_ may increase significantly at the same time. This scenario, in which *α* is fixed at 3, is considered in Table 3, too. Finally, it should be noted that, according to the recent literature, ^35^ NSE intensities for low *q* can be affected by additional effects. We have therefore checked the robustness of our treatment by leaving out *q* values below 0.047 Å^−1^ from the analysis. As seen in the Supporting Information (Table S1), the observed trends are conserved, with the only difference of overall slightly higher values of *κ*_1_.

In summary, the reasonable values obtained for the parameters *κ*_1_ and *α*, together with the observed trends regarding the influence of glycolipids, demonstrate that NSE data from polydisperse OLVs can be interpreted with the SANS-based procedure proposed here. The NSE data, in turn, revealed that saccharide-induced membrane coupling occurs on time scales relevant for membrane fluctuations.

## Conclusions

This study addressed the influence of adhesion-promoting glycolipids on the rigidity and mechanical coupling of adjacent lipid bilayers. Oligolamellar vesicles (OLVs) based on the phospholipid POPC, which contained defined fractions of such glycolipids, were investigated with small-angle neutron scattering (SANS) and neutron-spin-echo (NSE) spectroscopy. The SANS measurements revealed that the OLVs are characterized by substantial polydispersity not only in their radius but also in the number *n* of lamellae. We have therefore introduced a method that uses detailed knowledge of the distribution of *n* in order to extract meaningful information from NSE data. Key ingredient are the *q*-dependent intensity contributions from OLVs with different *n*. The intermediate scattering functions modeled in this way (Eq. 10) reproduce the experimental data with two parameters, the bending rigidity of individual bilayers, *κ*_1_, and an inter-bilayer coupling parameter *α*. By contrast, the experimental data cannot be modeled satisfactorily with the same effective bending rigidity for all *q* (Eq. 4). The analysis yielded *κ*_1_ ≈ 7 *k*_B_*T* and revealed that the presence of glycolipids indeed increases the mechanical inter-bilayer coupling, at least when plausibly assuming that small fractions of glycolipids do not dramatically increase the rigidity of individual bilayers. In future studies, additional reference measurements with small unilamellar vesicles (SUV) for each lipid composition may be helpful to disentangle *κ*_1_ and *α* even better.

## Materials and Methods

### Materials

1-palmitoyl-2-oleoyl-glycero-3-phosphocholine (POPC) as well as Digalactosyl-diacylglycerol and N-hexadecanoyl-lactosyl-ceramide with unsaturated alkyl chains (DGDG and LacCer, respectively) were purchased form Avanti Polar Lipids (Alabaster, Alabama, US). Chloroform (purity ≥ 99.9%), methanol (purity ≥ 99.9%), and ethanol (purity ≥ 99.8%) were purchased from Sigma-Aldrich and used without further purification. Double-deionized ultrapure water (resistivity = 18.2 MΩ.cm) was obtained from a water purification station (Purelab classic, Elga, Celle, Germany). All glassware and sample vials were either newly unsealed or washed with chloroform before use.

### Sample Preparation

Solutions of glycolipid-phospholipid mixtures were prepared by individually dissolving lipid powder at concentrations of 1 mg/mL in a mixture of chloroform and methanol (4:1 by volume) for glycolipids and pure chloroform for phospholipids, and then mixing them together to ensure homogeneous lipid mixtures. In this way five differently mixed lipid solutions were obtained: POPC with 10% DGDG-unsat, POPC with 20% DGDG-unsat, POPC with 10% LacCer-unsat, POPC with 20% LacCer-unsat, and pure POPC.

The lipid solutions were first dried in their glass vials under a N_2_-flow for several hours to remove solvent residues. Afterwards the vials were filled up with D_2_O to a concentration of 5 mg/mL and placed in a sonication bath for ten minutes to detach the lipids from the inner walls of the vials. Formation of OLVs from initially large multilamellar aggregates was achieved via mechanical extrusion through a porous membrane with an extruder (Avanti Polar Lipids), with a pore diameter of 400 nm by specification. The vesicle suspensions (3 mL per sample) were then transferred into newly opened Falcon tubes and from there later into SANS/NSE measurement cuvettes.

### Dynamic light scattering (DLS)

DLS was carried out with a Multiangle light scattering spectrometer (LS Instruments, Fribourg, Switzerland). The samples were diluted 1000 *×* to a concentration of 1 *µ*g/mL, so that the particle interactions and multiple scattering both become negligible. The cuvette was placed in an index-matching bath of decahydronaphthalene having a controlled temperature (20.0 °C) with a precision of *±*0.1 °C. The measurements were performed with a solid-state laser of wavelength of *λ* = 660 nm and a maximum power of 100 mW. The scattering signal, measured at up to 17 angles between 40° and 120° corresponding to 0.00065 Å^−1^ ≤ *q* ≤ 0.00165 Å^−1^, was detected with two avalanche photodiodes and was analyzed with a pseudo-cross-correlation in the LS-Instruments correlator. The measurement time for each angle was 30 s. From the auto-correlation data, the average radius *R*_avg_ and the polydispersity index (PDI = [*δR/R*_avg_]^2^) were extracted with a self-written script based on a third-order cumulant analysis, as established by Mailer et al. ^46^

### Small-angle neutron scattering (SANS)

#### Experiments

SANS measurements were performed on the instrument D22 at the Institut Laue-Langevin (Grenoble, France) using a wavelength of *λ* = 6 Å (with Δ*λ/λ* = 0.1), and sample-to-detector distances of 17.6 m and 5.6 m for the rear detector and 1.3 m for the front panel detector. In this way, the covered *q*-range was 0.004 Å^−1^ ≤ *q* ≤ 0.72 Å^−1^, where *q* = (4*π/λ*) sin *θ* and 2*θ* is the scattering angle.

#### Modeling

The contribution of each OLV to the SANS signal was calculated as the form factor intensity of a spherical object with the respective shell structure, which can be readily calculated by summing over the well-known spherical form factors of all sub-shells ^47,48^ with SLD difference Δ*ρ* = |*ρ*_W_ − *ρ*_B_|. The finite *q*-resolution of the experimental data was taken into account in the modeling process, by convolution with a Gaussian function of width *δq* specified by the instrument settings. The best-matching model parameters were then obtained via *χ*^2^ minimization. Estimates of the statistical parameter errors are valid only within the framework of a ”perfect model”, characterized by a reduced *χ*^2^ close to unity.^49^ For the parameters of the SANS fits, in view of significant additional contributions due to systematic errors, much larger error estimates are therefore provided. They approximately reflect the variation of the obtained parameters throughout the evolution and refinement of the model description, i.e., they reflect the robustness of the parameters with respect to the model, and we therefore consider them more meaningful.^30^

### Neutron-spin-echo (NSE) measurements

NSE measurements were performed on the instrument IN15 at the Institut Laue-Langevin (Grenoble, France) using wavelengths (maximum Fourier time in parentheses) of 6 Å (42 ns), 8 Å (99 ns), 10 Å (194 ns), and 12 Å (335 ns) at detector angles of 8°, 7.5°, 6°, and 4°, respectively, such that a *q*-range of 0.03 Å^−1^ ≤ *q* ≤ 0.17 Å^−1^ was covered. D_2_O was subtracted as background and the measurements were corrected for resolution effects using graphite.

## Supporting Information

Additional SANS data with fits; Non-exponential decay of *S*(*q, t*) with *t*; Effective Γ values for individual samples when modeled with Eq. 4; *q*-dependence of Γ when modeled with Eq. 4; SANS fit attempt with nominal lipid volume fraction; NSE results when omitting data at *q <* 0.047 Å^−1^ in the analysis Additional NSE data with fits based on Eq. 10.

## Acknowledgement

Allocation of beamtime by the ILL is gratefully acknowledged. The raw data of the experiments can be found at https://doi.ill.fr/10.5291/ILL-DATA.9-13-921. Help with the SANS measurements from S. Prévost and O. Matsarskaia is gratefully acknowledged.

