## Supporting Information for "Small-Angle and Quasi-Elastic Neutron Scattering from Polydisperse Oligolamellar Vesicles Containing Glycolipids"

### Additional SANS data with fits

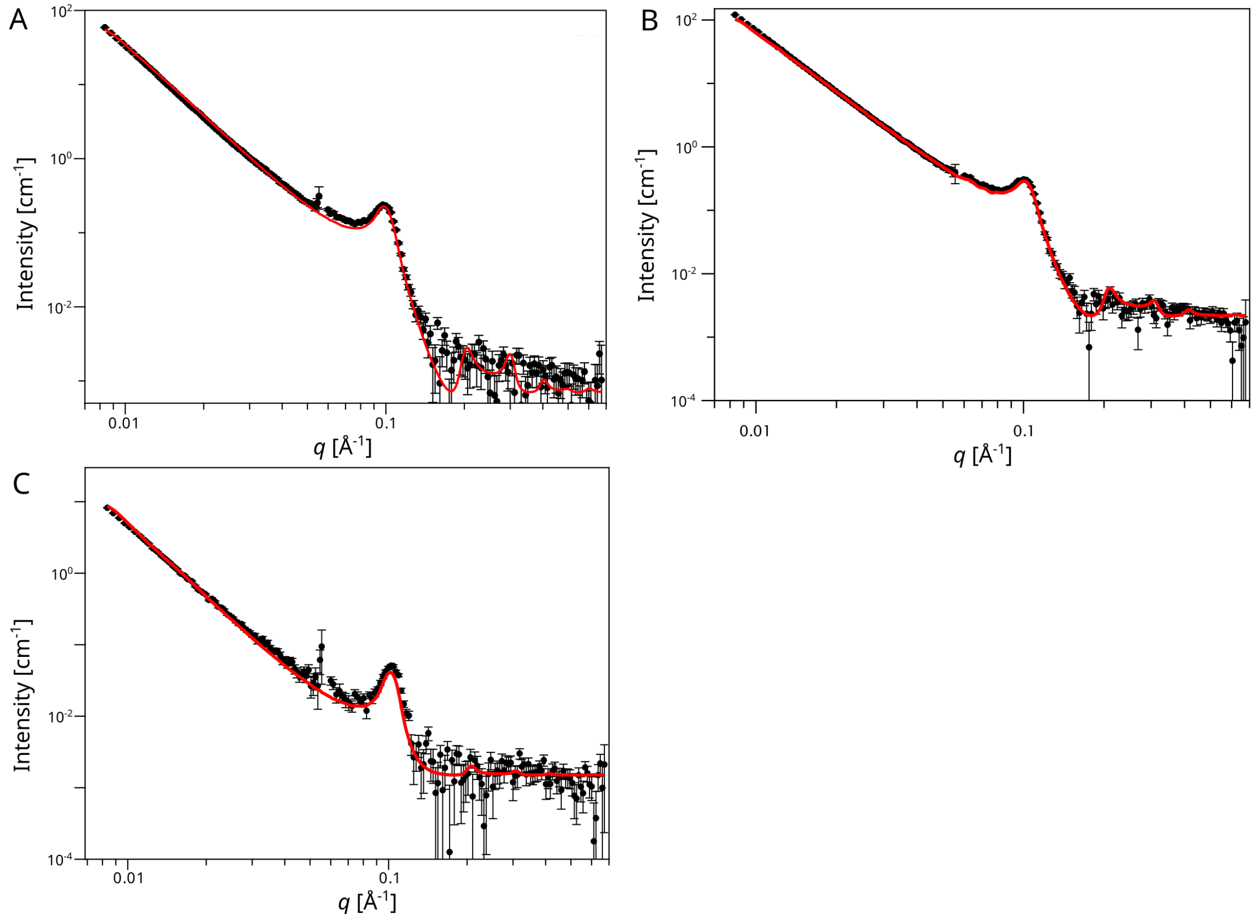


Figure S1: SANS curves after background subtraction, obtained with POPC OLVs (A), with POPC OLVs containing 10% DGDG (B), and with POPC OLVs containing 20% LacCer (C). Lines indicate fits to the data points.

### Non-exponential decay of $S(q,t)$ with $t$

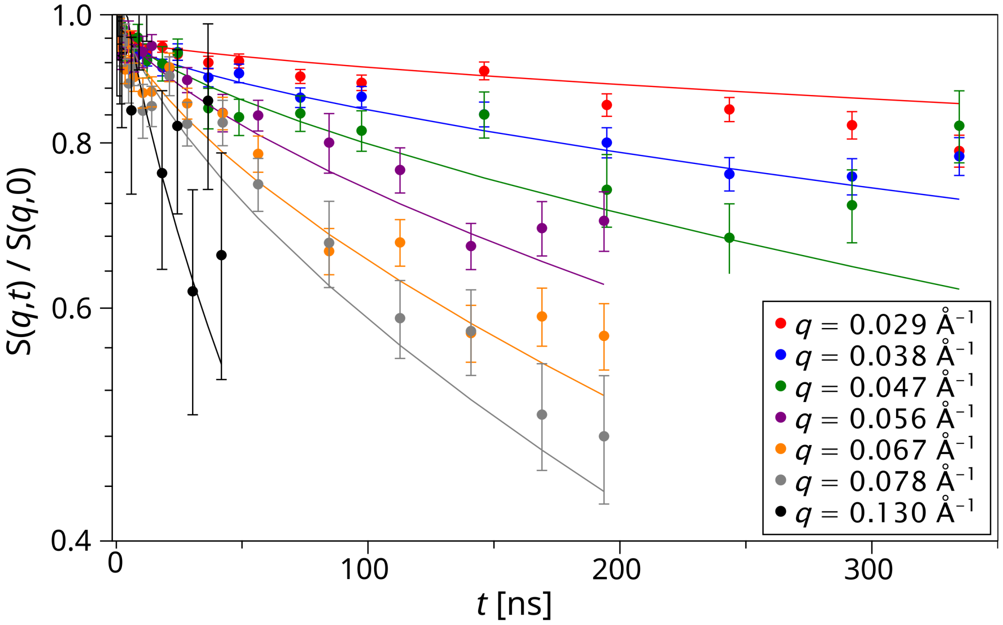


Figure S2: Semi-logarithmic representation of $S(q,t)$ for POPC OLVs containing 20% DGDG.

#### Effective $\Gamma$ values for individual samples when modeled with Eq. 4

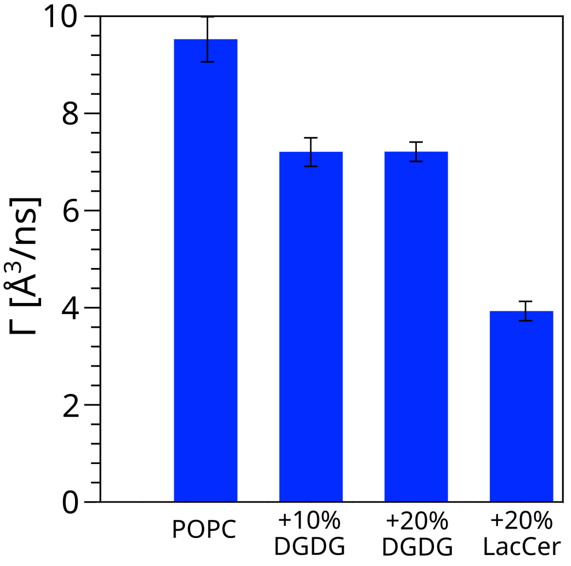


Figure S3: Effective $\Gamma$ values obtained for the individual samples when the whole data set $S(q,t)$ is fitted with Eq. 4. Error estimates correspond to the one-sigma confidence interval.

### $q$-dependence of $\Gamma$ when modeled with Eq. 4

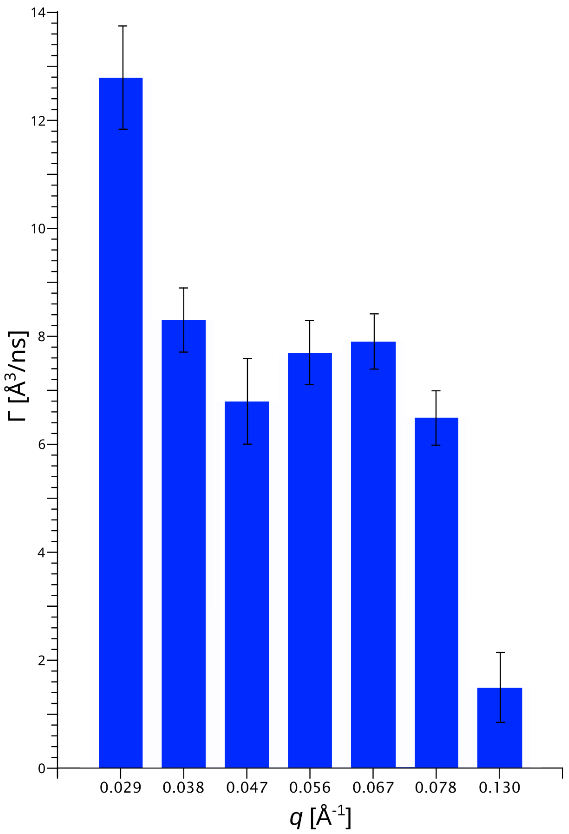


Figure S4: $\Gamma$ values obtained by fitting $S(q,t)$ independently for each $q$ value using Eq. 4. Data and fits are those shown in Fig. S2. Error bars correspond to the one-sigma confidence interval.

### SANS fit attempt with nominal lipid volume fraction

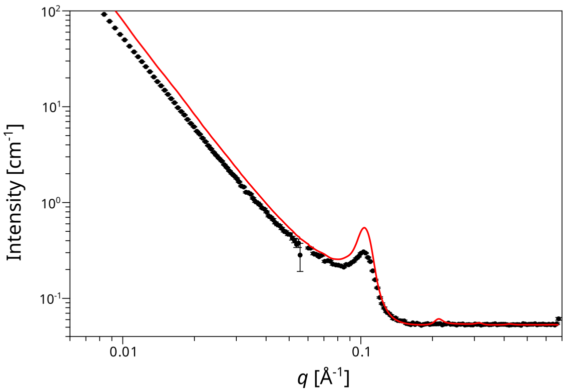


Figure S5: SANS curve after background subtraction, obtained with POPC OLVs containing 20% DGDG. The solid line indicates the attempt to fit the data with the nominal lipid volume fraction of 0.48 vol%.

### NSE results when omitting data at $q < 0.047$ Å$^{-1}$ in the analysis

Table S1: Alternative NSE results when omitting data at $q < 0.047$Å$^{-1}$.

| **OLV sample** | $\kappa_1$ [$k_B T$] (±1.0) | $\alpha$ (±0.5) |
|---|---|---|
| POPC | 14.2 | 1.0 (fixed) |
| | 11.3 | 2.0 (fixed) |
| +10%DGDG | 14.0 | 1.0 (fixed) |
| | 11.0 | 2.0 (fixed) |
| | 14.2 (fixed) | 1.0 |
| +20% DGDG | 13.7 | 1.0 (fixed) |
| | 11.6 | 2.0 (fixed) |
| | 14.2 (fixed) | 1.1 |
| +20% LacCer | 40.1 | 1.0 (fixed) |
| | 27.1 | 2.0 (fixed) |
| | 20.1 | 3.0 (fixed) |
| | 14.2 (fixed) | 5.2 |

### Additional NSE data with fits based on Eq. 10

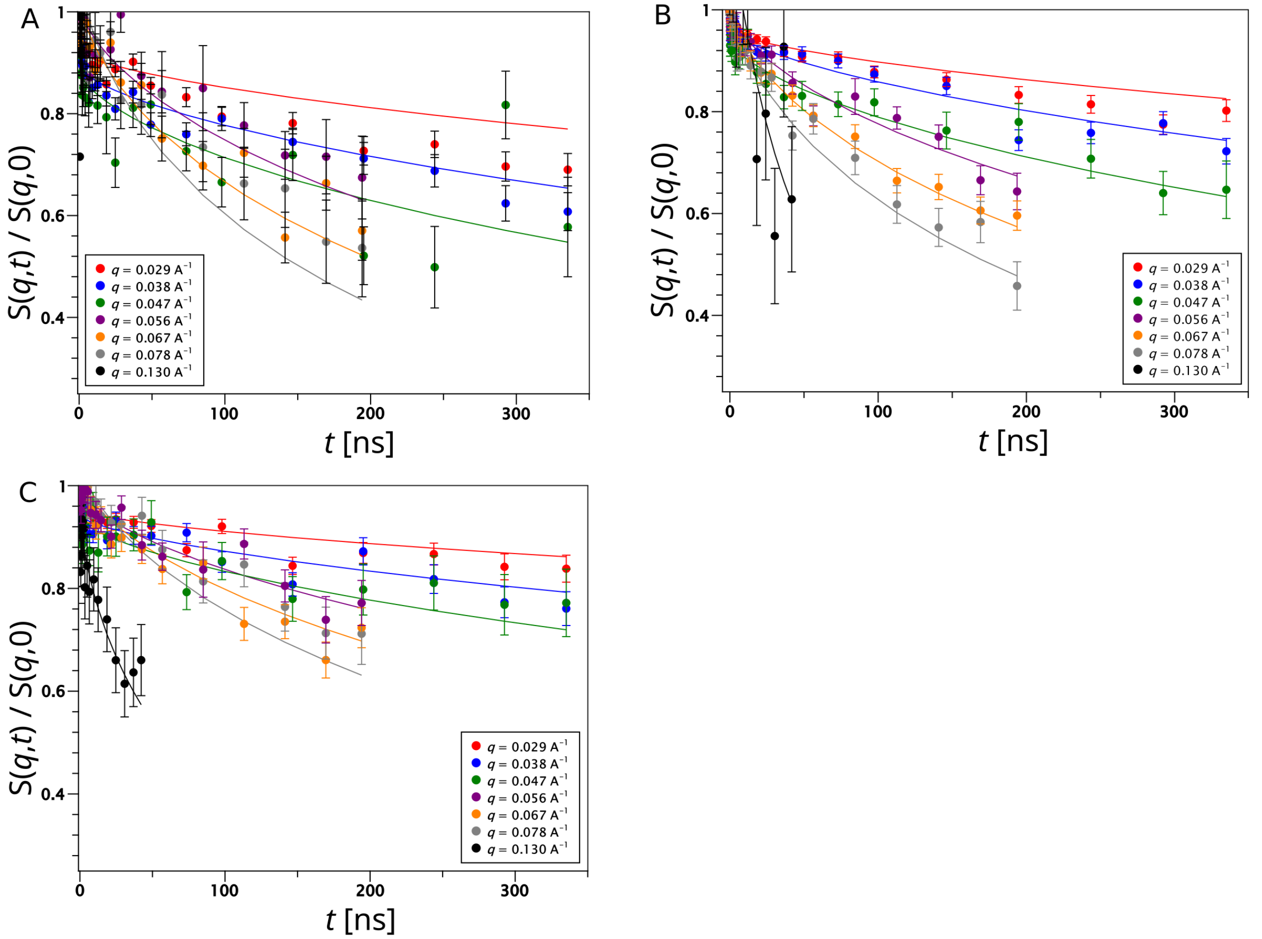


Figure S6: NSE data $S(q,t)$ obtained with POPC OLVs (A), with POPC OLVs containing 10% DGDG (B), and with POPC OLVs containing 20% LacCer (C). Lines indicate fits to the whole data sets using Eq. 10.
